# Mutation-related apparent myelin, not axon density, drives white matter differences in premanifest Huntington’s disease: Evidence from *in vivo* ultra-strong gradient MRI

**DOI:** 10.1101/2021.11.29.469517

**Authors:** Chiara Casella, Maxime Chamberland, Pedro Luque Laguna, Greg D. Parker, Anne E. Rosser, Elizabeth Coulthard, Hugh Rickards, Samuel C. Berry, Derek K. Jones, Claudia Metzler-Baddeley

## Abstract

White matter (WM) alterations have been observed early in Huntington’s disease (HD) progression but their role in the disease-pathophysiology remains unknown. We exploited ultra-strong-gradient MRI to tease apart contributions of myelin (with the magnetization transfer ratio), and axon density (with the restricted volume fraction from the Composite Hindered and Restricted Model of Diffusion) to WM differences between premanifest HD patients and age- and sex-matched controls. Diffusion tensor MRI (DT-MRI) measures were also assessed. We used tractometry to investigate region-specific changes across callosal segments with well-characterized early- and late-myelinating axonal populations, while brain-wise alterations were explored with tract-based cluster analysis (TBCA). Behavioural measures were included to explore disease-associated brain-function relationships. We detected lower myelin in the rostrum of patients (tractometry: p = 0.0343; TBCA: p = 0.030), but higher myelin in their splenium (p = 0.016). Importantly, patients’ myelin and mutation size were positively associated (all p-values < 0.01), indicating that increased myelination might be a direct result of the mutation. Finally, myelin was higher than controls in younger patients but lower in older patients (p = 0.003), suggesting detrimental effects of increased myelination later in the course of the disease. Higher FR in patients’ left cortico-spinal tract (CST) (p = 0.03) was detected, and was found to be positively associated with MTR in the posterior callosum (p = 0.033), possibly suggesting compensation to myelin alterations. This comprehensive, ultra-strong gradient MRI investigation provides novel evidence of CAG-driven myelin alterations in premanifest HD which may reflect neurodevelopmental, rather than neurodegenerative disease-associated changes.

## Introduction

Huntington’s disease (HD), a neurodegenerative disorder leading to devastating cognitive, psychiatric and motor symptoms, cannot currently be cured, and a research priority is to increase understanding of its pathogenesis. Subtle and progressive white matter (WM) alterations have been observed early in HD progression ^1–7^, but their aetiology and role remain unclear. Therefore, with the present study we aimed to disentangle the contribution of changes in axon microstructure *versus* changes in myelin to WM pathology in premanifest HD. Crucially, we exploited the very latest-in ultra-strong magnetic field gradient technology ^8,9^ to achieve high-b-values and increased restriction of water diffusion. In turn, this afforded an enhanced differential attenuation of intra- and extra-axonal MRI signals, while maintaining sufficient signal-to-noise ratio (SNR), and thus allowed to better tease apart the contribution of different sub-compartments of WM microstructure ^10–12^.

More specifically, we assessed WM microstructure in premanifest patients by combining fractional anisotropy (FA), axial diffusivity (AD) and radial diffusivity (RD) from diffusion tensor (DT)-MRI ^13^, with the magnetization transfer ratio (MTR) from magnetization transfer imaging (MTI) as a proxy measure of myelin, and the restricted diffusion signal fraction (FR) from the Composite Hindered and Restricted Model of Diffusion (CHARMED) ^14^ as a proxy measure of axon density ^15^. Alterations in microstructural metrics were assessed using two analytical pipelines: i. a tractometry approach ^16–18^ to assess tract-specific changes across the corpus callosum (CC), and ii. a whole-brain approach ^19^ to further explore abnormalities associated with the premanifest disease stage.

The CC is the brain’s largest WM tract and its fibres vary in size and age of myelination, with larger, early myelinating fibres seen in posterior portions, and smaller, later-myelinating fibres found in anterior callosal regions ^20^. Thus, characterising WM microstructure across this tract might provide insight into regional differences in the impact of HD on WM, and elucidate disease-related pathological processes in the context of the Demyelination Hypothesis ^21^. This suggests that mutant huntingtin (*mHTT)* leads to premature myelin breakdown and has been given support by several studies demonstrating alteration in myelin-associated biological processes at the cellular and molecular level in the HD brain ^7,22–28^. Specifically, the Demyelination Hypothesis proposes myelin impairment to begin from early-myelinating caudate and putamen striatum structures and then spread in a bilateral and symmetric pattern to other early-myelinating regions. Thus, in the context of the present study, the Demyelination Hypothesis would predict more dominant microstructural changes in posterior relative to anterior callosal subregions, as the former myelinate earlier.

Following evidence that WM volume loss in HD extends beyond the CC ^2,29–35^, and the concept of compensatory networks in response to neurodegeneration ^36^, we supplemented the tractometry analysis with a novel exploratory, whole-brain analysis, called Tract-Based Cluster Analysis (TBCA)^19^ to assess brain-wise group microstructural differences. TBCA uses the rich anatomical information from whole-brain tractography reconstructions, to inform the cluster-level inference analysis of voxel-based images, and provides the anatomical specificity required to disentangle distinct clusters belonging to different anatomical tracts ^19^.

Finally, the evidence of cognitive and behavioral impairments in premanifest patients ^2,35,37^ across attention, working memory, processing speed, psychomotor functions, episodic memory, emotion processing, sensory-perceptual functions, and executive functions ^2,38–43^, and their significant impact on everyday functional decline ^44–46^, stress the importance of understanding how these symptoms may relate to pathological neural changes, such as alterations in WM microstructure. For this purpose, we derived a composite cognitive score using principal component analysis (PCA) to capture variability in patients’ cognitive performance, and used it for the analysis of correlations between differences in cognition and WM microstructure.

## Materials and methods

### Participants

25 individuals with premanifest HD and 25 age- and sex-matched healthy controls were recruited, with ethical approval from the local National Health Service (NHS) Research Ethics Committee (Wales REC 5 18/WA/0172) and by the Cardiff University School of Psychology Ethics Committee. All participants provided written informed consent prior to taking part in the study.

Patients were recruited from the Cardiff HD Research and Management clinic, Bristol Brain Centre at Southmead Hospital, and the HD clinic at the Birmingham and Solihull NHS Trust. Healthy controls were recruited from Cardiff University and the School of Psychology community panel. Participants were recruited if eligible for MRI scanning. Control participants were excluded if they had a history of neurological or psychiatric conditions, and patients if they had a history of any other neurological condition.

22 of the HD patients had pen-and-paper cognitive task data available from their most recent participation in the ENROLL-HD study (NCT01574053, https://enroll-hd.org). The progression of symptoms in ENROLL-HD participants is monitored longitudinally, and one of the optional components within the study is the giving of permission by participants for their coded data to be accessed by researchers in the field. As such, a full clinical dataset including full medical and medication history is available for each research participant and some of these data were used in this study.

One control subject was excluded from the tractometry analysis because of poor callosal segmentation. Therefore, data from 25 patients and 24 healthy controls were used for callosal tractometry analysis. As this did not impact TBCA, a sample of 25 patients and 25 controls was analysed. Table 1 provides a summary of participants’ demographic and clinical background information. Performance in the Montreal Cognitive Assessment (MoCA) ^47^ and in the Test of Premorbid Functioning - UK Version (TOPF-UK) ^48^ is reported for patients and controls. The Unified Huntington Disease Rating Scale (UHDRS) total motor score (TMS), total functional capacity (TFC), diagnostic confidence level (DCL) and CAG repeat size obtained from the ENROLL-HD database are also reported for patients.

**Table 1.**
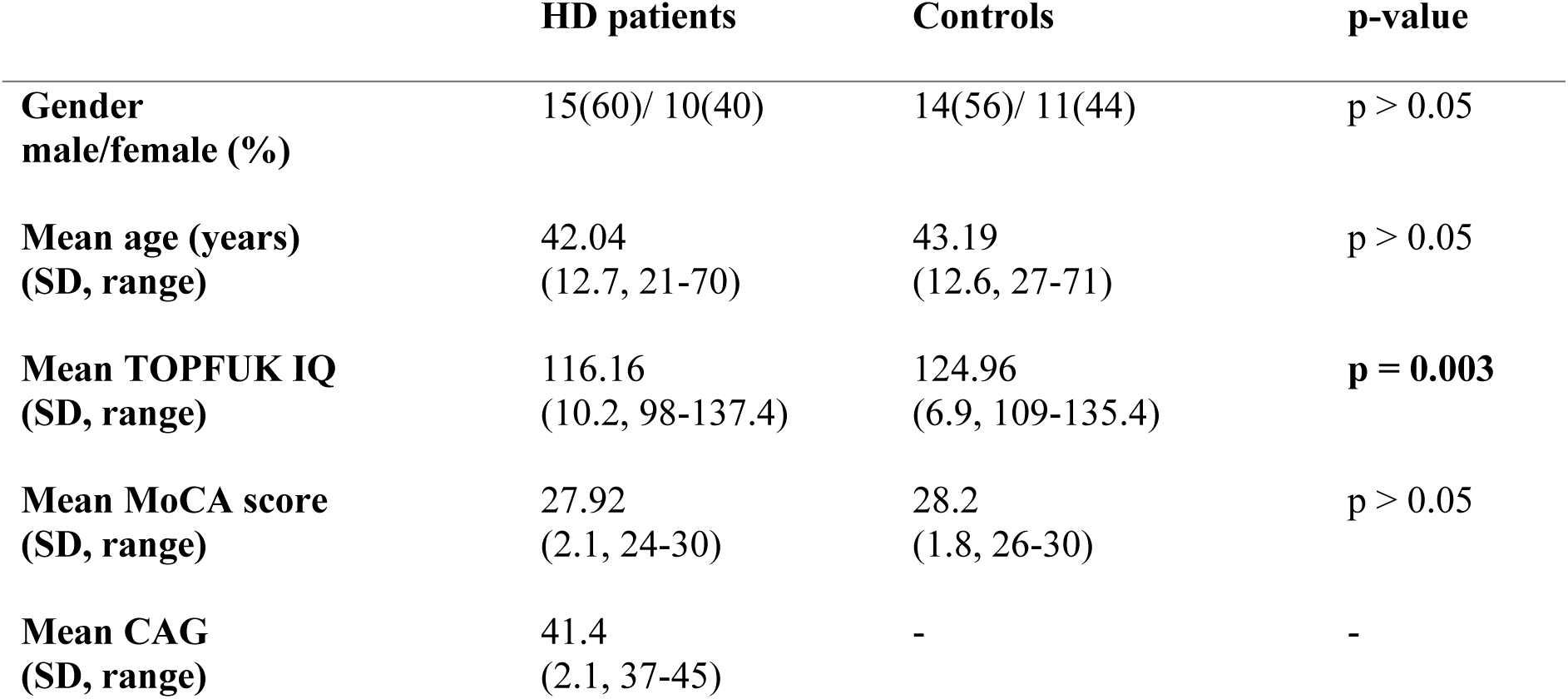
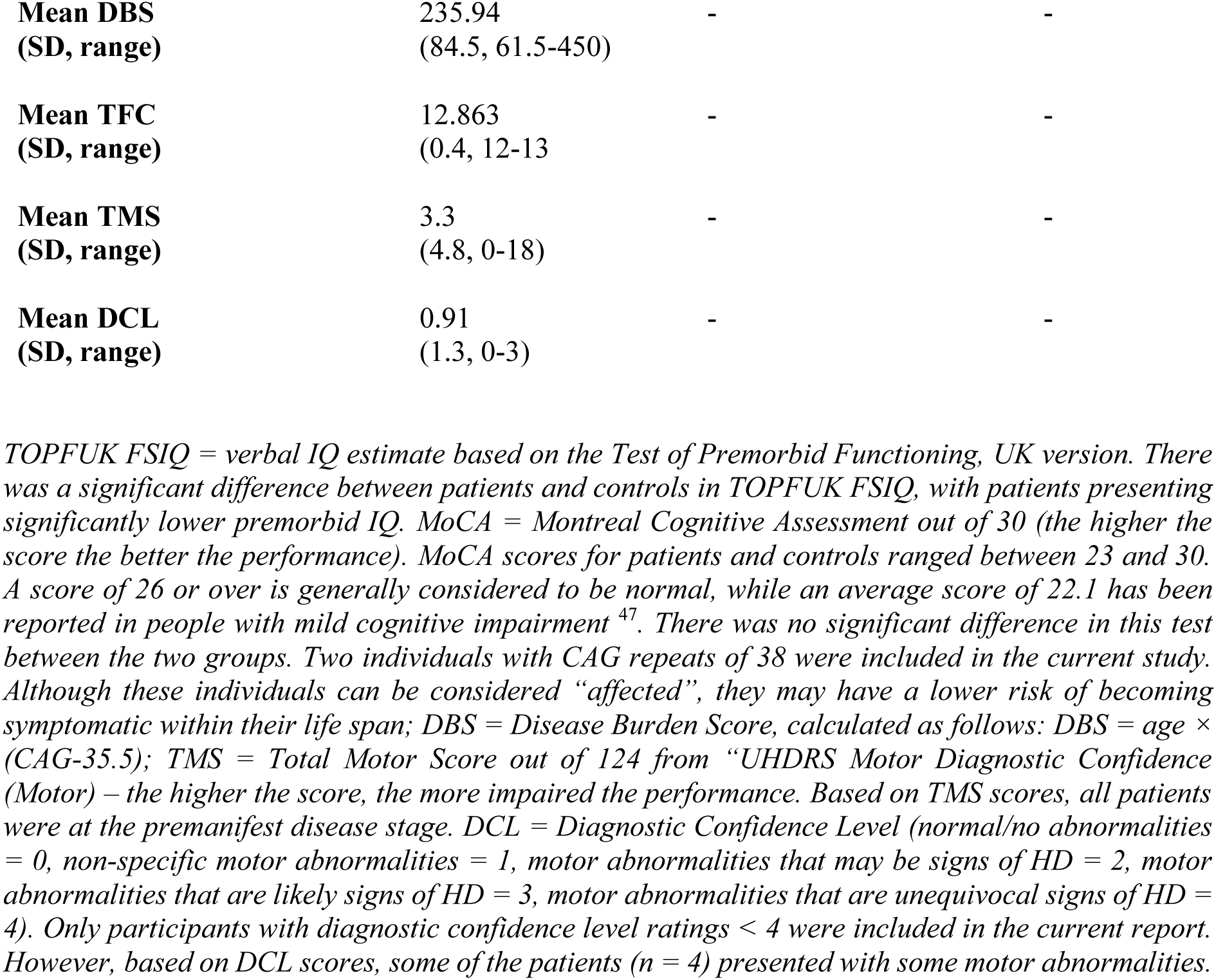
Summary of participants’ demographic and clinical background information.

|  | **HD patients** | **Controls** | **p-value** |
| --- | --- | --- | --- |
| **Gender**<br>**male/female (%)** | 15(60)/ 10(40) | 14(56)/ 11(44) | p > 0.05 |
| **Mean age (years)**<br>**(SD, range)** | 42.04<br>(12.7, 21-70) | 43.19<br>(12.6, 27-71) | p > 0.05 |
| **Mean TOPFUK IQ**<br>**(SD, range)** | 116.16<br>(10.2, 98-137.4) | 124.96<br>(6.9, 109-135.4) | **p = 0.003** |
| **Mean MoCA score**<br>**(SD, range)** | 27.92<br>(2.1, 24-30) | 28.2<br>(1.8, 26-30) | p > 0.05 |
| **Mean CAG**<br>**(SD, range)** | 41.4<br>(2.1, 37-45) | - | - |
| **Mean DBS**<br>**(SD, range)** | 235.94<br>(84.5, 61.5-450) | - | - |
| **Mean TFC**<br>**(SD, range)** | 12.863<br>(0.4, 12-13) | - | - |
| **Mean TMS**<br>**(SD, range)** | 3.3<br>(4.8, 0-18) | - | - |
| **Mean DCL**<br>**(SD, range)** | 0.91<br>(1.3, 0-3) | - | - |

### Data acquisition

#### Assessment of disease-related brain-function relationships

A composite cognitive score was computed by combining cognitive data available for patients on the ENROLL-HD database (providing these had been obtained within a 3-month time window from their participation in the present study), with data acquired during the study. This was done in order to reduce patient burden associated with study participation. Table 2 provides details on the administered tests, the cognitive domains they assess, and the outcome variables measured.

**Table 2.**
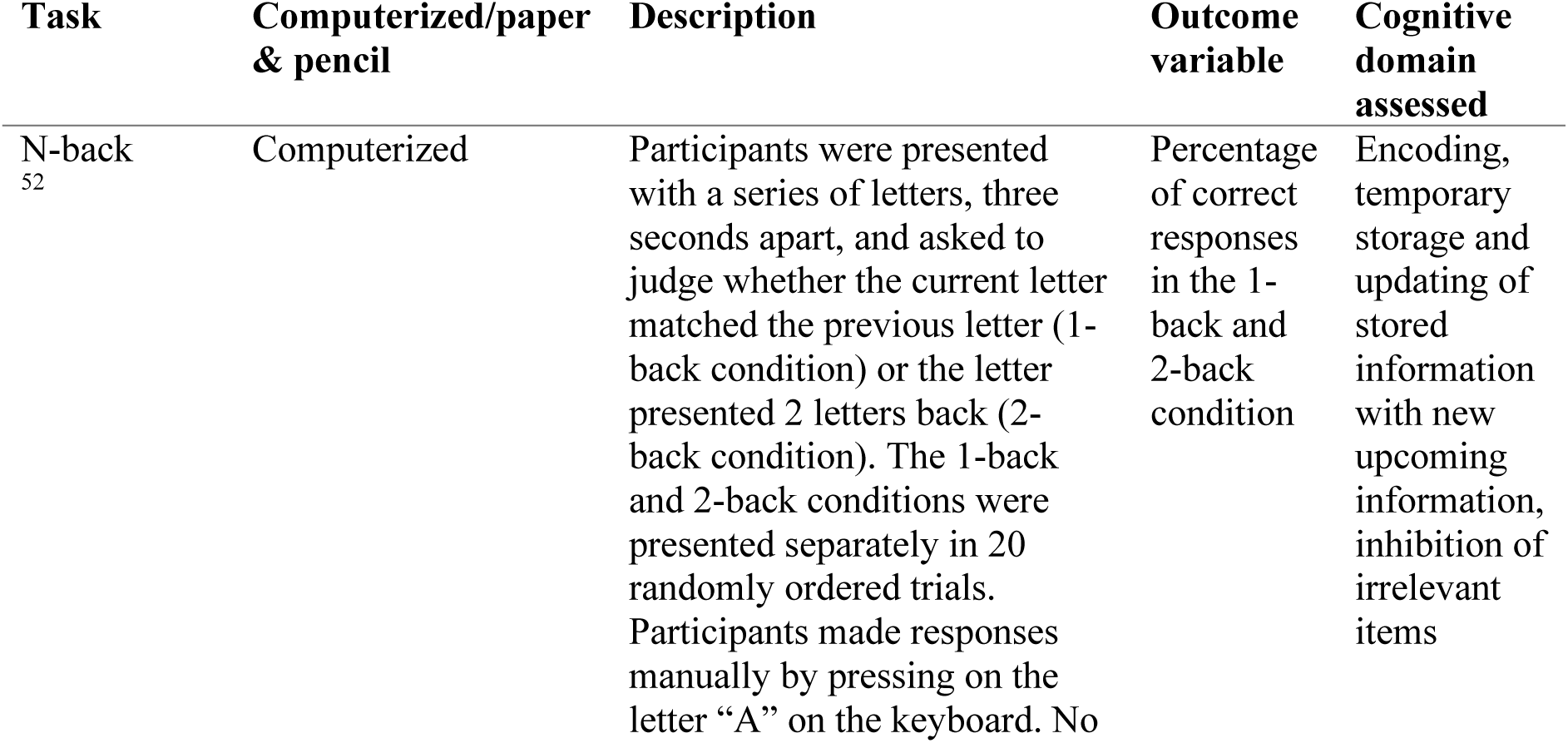
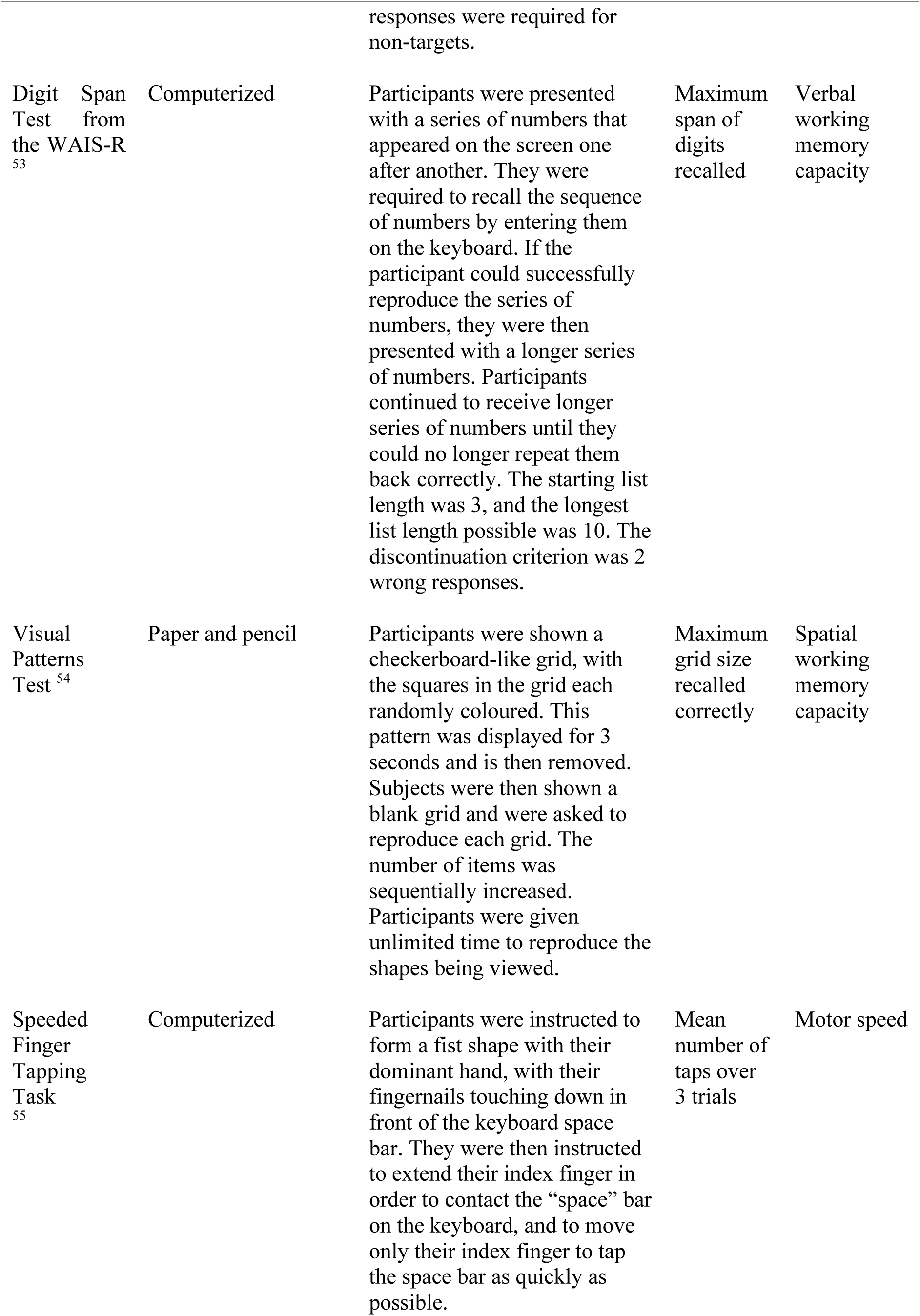
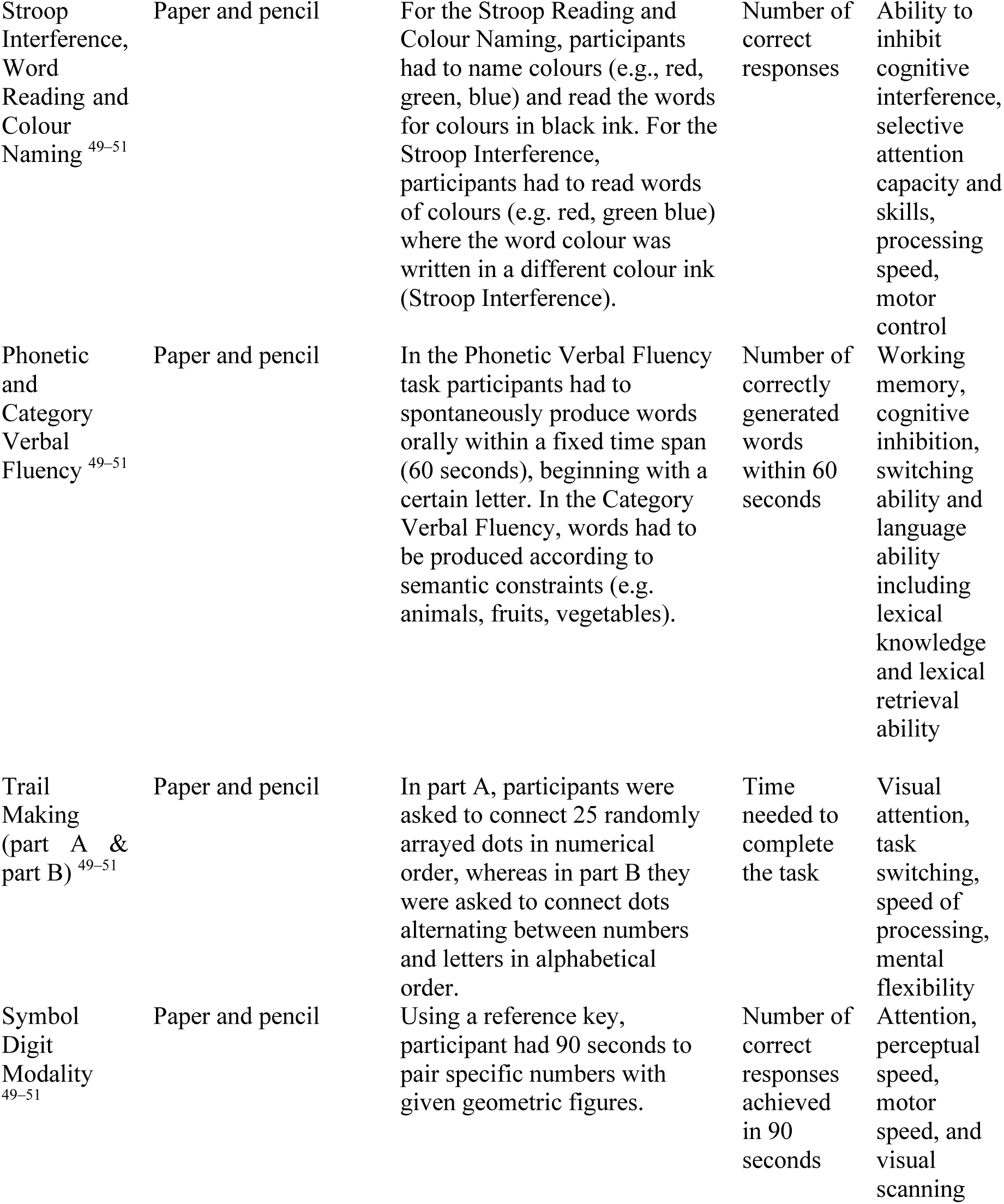
Cognitive outcome variables employed to create a composite cognitive score to assess disease-related brain-function relationships. Tasks descriptions are provided, outcome variables and cognitive domains assessed are summarized.

Briefly, data from the ENROLL-HD database concerned performance in the Phonetic Verbal Fluency Test, the Categorical Verbal Fluency Test, the Symbol Digit Modality Test, the Stroop Colour Reading and Word Reading Test, the Stroop Interference Test and the Trail Making Test ^49–51^– please see http://www.enroll-hd.org for the detailed study protocol.

On the other hand, performance in the N-back Task ^52^, the Forward Digit Span Test adapted from the Wechsler Adult Intelligence Scale-Revised (WAIS-R) ^53^, the Visual Patterns Test ^54^ and the Speeded Finger Tapping Task ^55^ was assessed as part of the present study. Cognitive testing was performed prior to MRI scanning and lasted approximately 60 minutes. Tasks were administered either as paper and pencil tests or by using a computerized version provided by the Psychology Experiment Building Language (PEBL) test battery ^56^.

As each task yields several outcome variables, the following strategy was employed: (1) for standardized clinical tests, metrics known to have the best sensitivity and measurement characteristic were selected, e.g. correctly-generated responses instead of error scores ^57^; (2) for tests with multiple conceptually distinct outcome measures, variables that represented each component were included, e.g., for the N-back Task, the number of correct responses from the 1-back and the 2-back condition; and (3) where necessary, variables were excluded from the assessment, e.g. when these presented lots of missing cases. This approach led to 13 cognitive outcome measures (Table 2).

#### MRI data acquisition

MRI data were acquired on a 3 Tesla Siemens Connectom system with ultra-strong (300 mT/m) gradients. Each MRI session lasted 1 hour, and comprised: a T_1_-weighted MPRAGE; a multi-shell dMRI acquisition [δ/Δ: 7/24 ms; b-values: 0 (14 volumes, interleaved), 500 (30 directions), 1200 (30 directions), 2400 (60 directions), 4000 (60 directions), and 6000 (60 directions) s/mm^2^ ^59^. Data were acquired in an anterior-posterior phase-encoding direction, with one additional posterior-to-anterior volume]; and a magnetization transfer acquisition [turbo factor: 4; radial reordering; non-selective excitation; MT contrast was achieved by the application of a 15.36 ms radio-frequency saturation pulse, with an equivalent flip angle of 333° applied at a frequency of 1.2 kHz below the water resonance. Two identical sets of images with different contrasts (one acquired with and one acquired without MT saturation pulses) were obtained]. Table 3 provides more details on the acquisition parameters.

**Table 3.** Scan parameters.

|  | **T<sub>1</sub>-w** | **DTI** | **CHARMED** | **MT** |
| --- | --- | --- | --- | --- |
| Pulse sequence | MPRAGE | SE\EPI | SE\EPI | Turbo FLASH |
| Matrix size | 256×256 | 495×495 | 495×495 | 128 × 128 ×104 |
| FoV (mm) | 256 | 990 | 990 | 220 × 220 × 179 |
| Slice thickness (mm) | 1 | 2 | 2 | 1.72 |
| TE,TR (ms) | 2, 2300 | 59, 3000 | 59, 3000 | 2.1, 60 |
| Off-resonance pulses (Hz/°) | - | - | - | 1200/333 |
| Flip angles (°) | 9 | 9 | 9 | 5 |

*All sequences were acquired at 3 Tesla with ultra-strong gradients. For each of the sequences, the main acquisition parameters are provided. T<sub>1</sub>-w: T<sub>1</sub>-weighted; MT: magnetization transfer; MPRAGE: Magnetization prepared - rapid gradient echo; SE: spin-echo; EPI: echo-planar imaging; FoV: field of view; TE: echo time; TR: repetition time.*

### Image processing

All images were skull-stripped in native space using FSL BET ^60^.

#### Diffusion data: FA, RD, AD, MD and FR maps

Pre-processing of diffusion data was carried out using FMRIB Sofware Library (FSL) ^60^, MRtrix3 ^61^, and Advanced Normalization Tools (ANTs) ^62^. These steps included: denoising ^63^, slice-wise outlier detection (SOLID) ^64^, and correction for drift ^65^; motion, eddy, and susceptibility-induced distortions ^66,67^; Gibbs ringing ^68^; bias field ^69^; and gradient non-linearities ^70,71^.

Diffusion tensors were estimated using linearly-weighted least squares regression (for b < 1200 s/mm^2^ data) providing the following quantitative scalar measures: FA, AD and RD. The diffusion tensor was fitted to data between b = 500 s/mm^2^ and b = 1200 s/mm^2^ in order to reduce cerebrospinal fluid based partial volume artefacts in the DTI metrics. The CHARMED data were corrected for motion and distortion artefacts ^72^, before computing FR maps ^14^ using in-house software coded in MATLAB (The MathWorks, Natick, MA).

#### Magnetization transfer: MTR maps

MT- and non-MT-weighted images were corrected for Gibbs ringing ^68^. ANTS ^62^ was first used to nonlinearly register the MPRAGE images to the b = 0 s/mm^2^ images. Then MT- and non-MT weighted images were linearly warped to the registered MPRAGE images using an affine (12 degrees of freedom) technique based on mutual information, with the FMRIB’s Linear Image Registration Tool (FLIRT)^73^. All registrations were visually inspected for accuracy. Finally, MTR maps were calculated according to: MTR = [(S^0^-S *^MT^*)/S^0^] × 100, whereby S^0^ represents the signal without the off-resonance pulse and S*^MT^* represents the signal with the off-resonance pulse.

#### Tractography of the CC

Automated WM tract segmentation of the CC was performed using TractSeg ^74^ and multi-shell constrained spherical deconvolution (MSMT-CSD) ^75^. Specifically, seven portions of the CC were delineated [1=rostrum, 2=genu, 3=rostral body, 4=anterior midbody, 5=posterior midbody; 6=isthmus, 7=splenium] (Fig. 1). For each segment, 2000 streamlines were generated.

**Figure 1.**
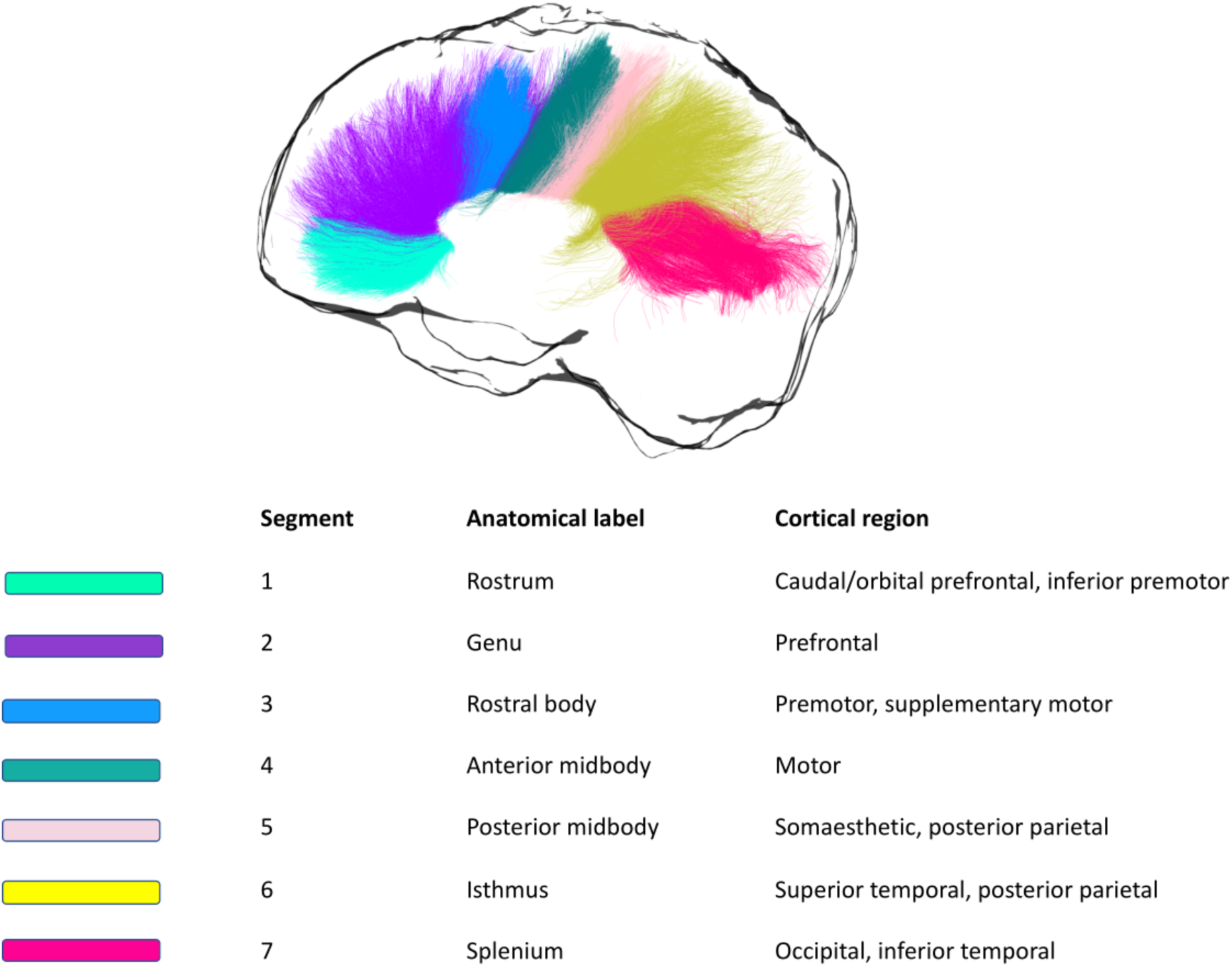
Callosal segmentation. For each segment, the corresponding anatomical label is reported, together with the cortical area it connects to.

### Statistical analysis

Analyses were performed in RStudio ^76^, MATLAB (The MathWorks, Natick, MA), SPSS ^77^, the PROCESS computational tool for mediation analysis ^78^, FSL ^60^, and the Statistical Non-Parametric Mapping (SnPM) software ^79^. Outliers were first identified by examining box-and-whisker plots for each dependent variable, for controls and patients separately. Outliers that were ± 3 standard deviations from the mean were removed.

#### Assessment of disease-related brain-function relationships

PCA of the cognitive data was performed on the slopes from patients, to best capture heterogeneity within this population. The first principal component (PC) only was extracted, to increase experimental power and reduce the number of multiple comparisons ^80^.

First, the Bartlett’s test of sphericity and the Kaiser-Meyer-Olkin (KMO) test were used to confirm that the data were suited for PCA [KMO = 0.54, χ2 (78) = 156.5, p <0 .001]. The PCA was run using centred, standardized versions of the patients’ cognitive outcome scores. Orthogonal Varimax rotation was used to maximize the factor loadings. Regression values from each component were used as composite cognitive scores for each patient.

#### Tractometry of the CC

Microstructure differences were assessed in the seven callosal segments. By taking each quantitative metric map, samples of each metric were obtained at each vertex of the reconstructed segments, and segment-specific medians were derived for FA, AD, RD, FR and MTR in MRtrix3 ^61^. Next, the overall mean was calculated, so that each dataset comprised *m* = 5 MRI-derived measures, mapped along s = 7 callosal segments.

#### Reduction of MRI data dimensionality with PCA

PCA was also employed to reduce the complexity of the callosal microstructure data ^81^. Centred, standardized versions of MRI measures on both groups combined were used ^82^. Specifically, the PCA was calculated for FA, FR, RD, AD and MTR, after checking that the data was suited for this analysis [KMO = 0.65, χ^2^ (6) = 1077.231, p < 0.001]. PCA was applied to the concatenated set of segments across subjects ^83,84^. The number of principal components was extracted based on: 1) their interpretability ^85^; 2) the Kaiser criterion of including all components with an eigenvalue greater than 1. Regression values from each component for each participant were used in the following analyses.

#### Investigation of group differences in callosal microstructure

To assess group differences in callosal microstructure, analyses of covariance (ANCOVAs) were run on the extracted regression values from each component for each participant. Group and segment were used as independent variables because of a particular interest in understanding the interaction between group effects on different callosal segments. The correlation of microstructure outcome measures across patients and controls, with age, ICV and TOPF-UK FSIQ was tested to decide if these variables should be included as covariates in the analysis. Pearson’s correlation coefficients greater than 0.3 were treated as indicative of a moderate relationship. For every ANCOVA, analysis assumptions were first tested.

#### Assessment of disease-related brain-function relationships

Spearman correlations were run in the patient group for:

i. WM components showing a significant group effect and composite cognitive scores;
ii. WM components showing a significant group effect and CAG repeat length;
iii. WM components showing a significant group effect and disease burden score (DBS), calculated as follows: DBS = age × (CAG-35.5).

Within each group of correlations, multiple comparison correction was carried out with Bonferroni with a family-wise alpha level of 5% (two-tailed). Whenever a significant association was detected, this was further explored with partial correlations, partialling out ICV and DBS. The latter was done to assess associations independently of disease progression.

#### TBCA assessment of brain-wise group differences in WM microstructure

TBCA ^19^ was applied to assess group differences in FA, RD, AD, FR and MTR. This method is based on the novel concept of ‘hypervoxel’, which extends standard 3D voxels with extra dimensions to encode geometrical and topological information about the streamlines that intersect each voxel.

All images were first non-linearly normalised to the FMRIB58_FA template (1 × 1 × 1 mm isotropic) using the tbss_2_reg script ^86^. Next, statistics maps were produced based on the voxel-level analysis of the data by using a non-parametric approach based on a permutation test strategy ^87^. The statistic maps were then thresholded by a value of p = 0.01, and the suprathreshold voxel-level statistic results were projected onto an hypervoxel template built on whole-brain tractography data from 20 healthy subjects. Two hypervoxels were defined as belonging to the same cluster if they were either adjacent or connected within the hypervoxel template (i.e. if they shared a common streamline) ^19^. Finally, the mass of each cluster ^88^ was computed and their corresponding statistical significance calculated based on the same permutation tests used for the voxel-level inference. Explanatory variables (EVs) in the permutation tests included age and gender and the effect of group was explored whilst regressing the other EVs. Clusters with a family-wise error (FWE)-corrected ^79^ p-value below 0.05 were considered statistically significant. A schematic representation of the TBCA pipeline can be found in Fig. 2.

**Figure 2.**
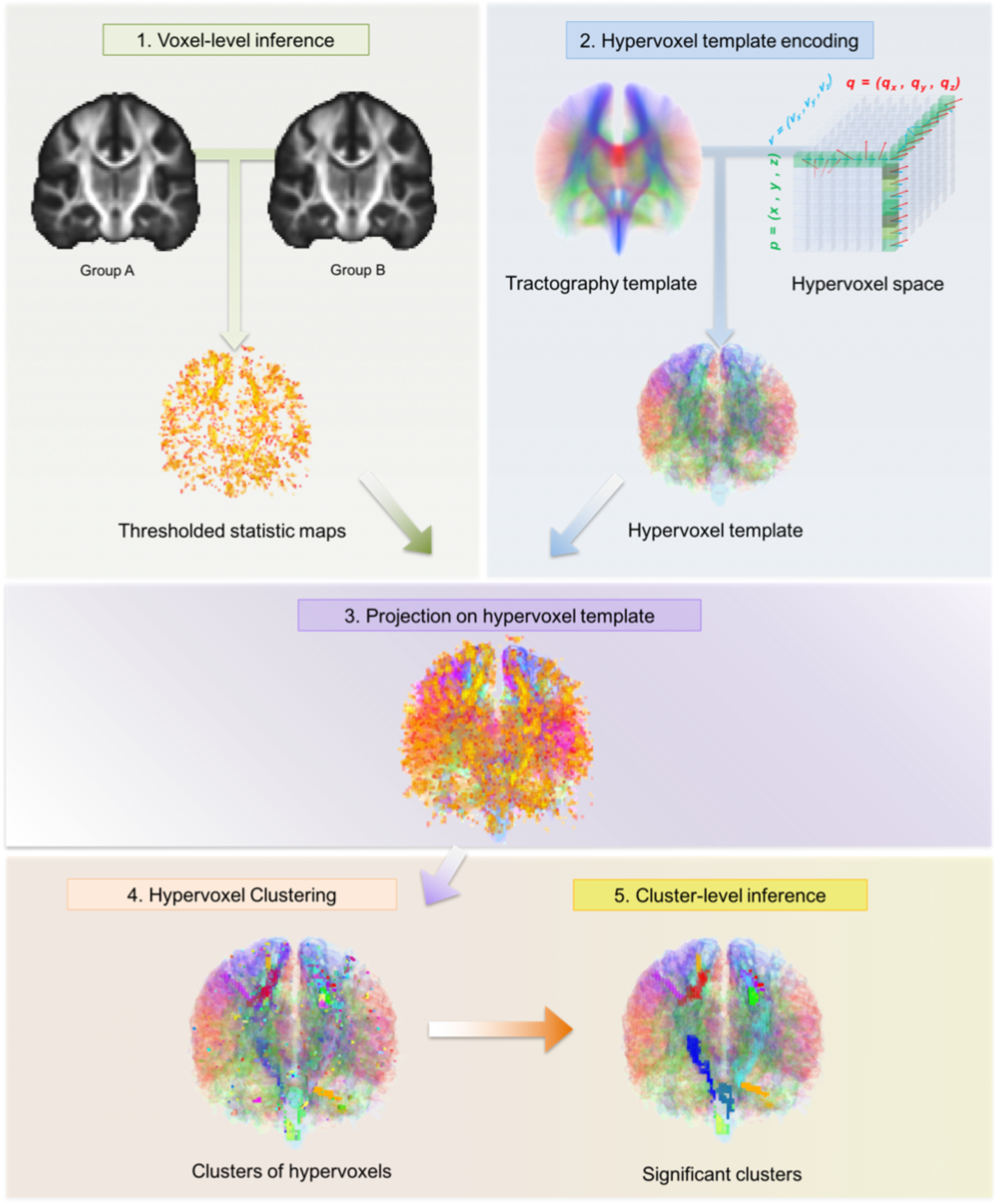
The TBCA analysis pipeline. After all images have been normalized to a common anatomical space, statistics maps are produced based on the voxel-level analysis of the data; this is done by using a non-parametric approach based on a permutation test strategy ^87^. The statistic maps are thresholded by a value of p = 0.01. Next, the significant voxel level statistic results are projected on a hypervoxel template. Finally, significant clusters of hypervoxels are identified.

Whenever significant clusters were detected for a specific metric, these were extracted, summed and binarized to form an ROI mask. The mask was then projected onto each map in MNI space. The mean value for that metric was calculated in the ROI with FSL ^60^, and used to run Spearman correlations between the WM metrics showing significant clusters. Multiple comparison correction was carried out with the Bonferroni correction with a family-wise alpha level of 5% (two-tailed).

## Results

### Composite cognitive score in the patient sample

As shown in Fig. 3, the first principal component (PC) accounted for 38.7% of the total variance in the cognitive data. Component loadings of >= 0.5 were considered as significant ^89^. Thus, this component reflected general executive functioning with loadings on distractor suppression (Stroop task), attention switching (Trail Making), updating (N-back), category fluency and motor speed.

**Figure 3.**
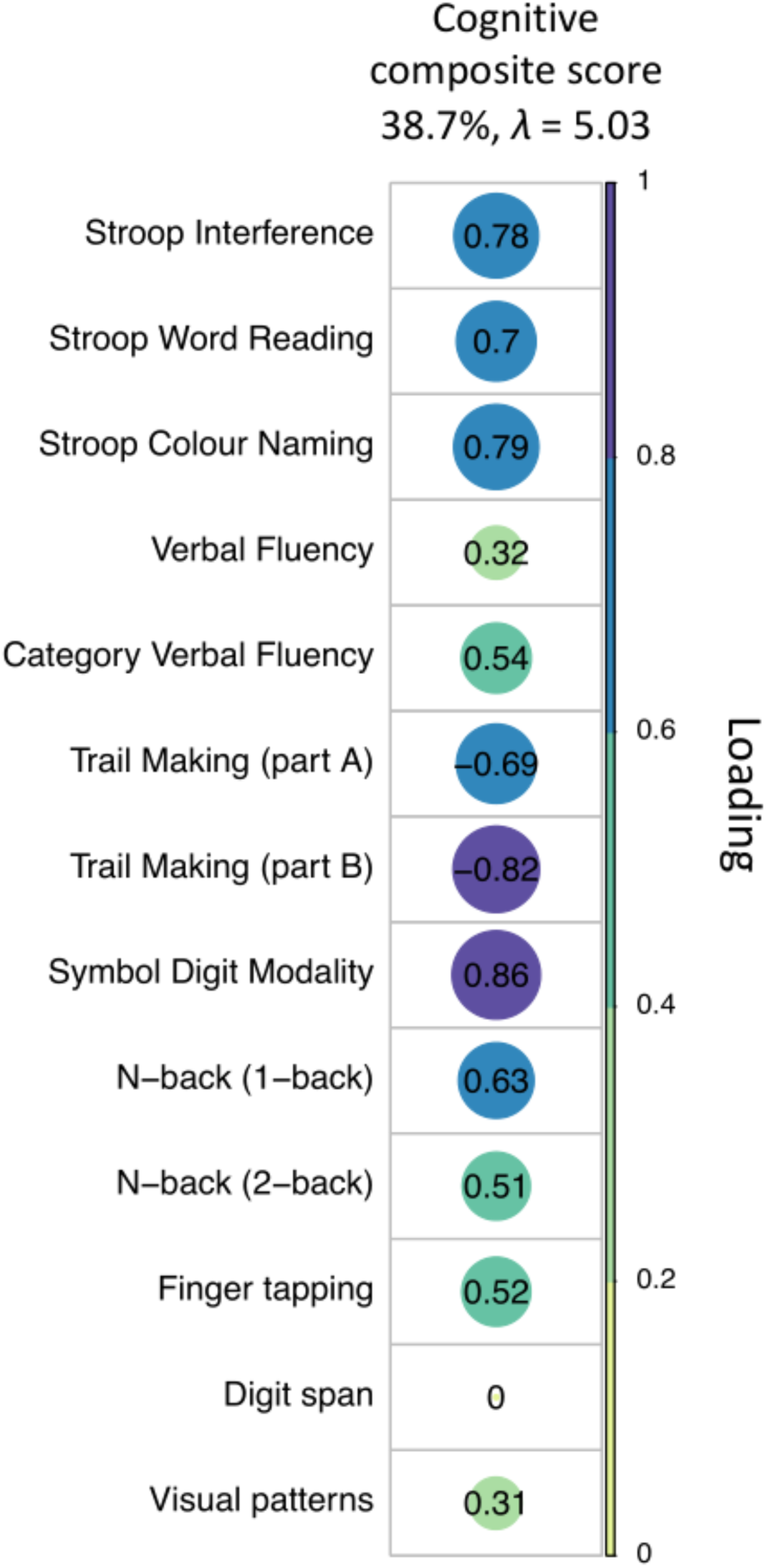
PCA of the cognitive data with varimax rotation. Plot summarizing how each variable is accounted for in the extracted PC. The absolute correlation coefficient is plotted. Color intensity and the size of the circles are proportional to the loading. This PC accounted for 38.7% of the total variance and included measures from all test domains, except for the digit span. Four patients were excluded from the PCA because of missing data. The final sample size for the PCA was n=21 patients.

#### Reduction of MRI data dimensionality with PCA

Over 80% of the variability in the microstructure data was accounted for by the first two PCs (PC1, 58.1%, *λ* = 2.90; PC2, 22.6%, *λ* = 1.13). As shown in Fig. 4, the first PC loaded positively on FA, FR, and AD, and negatively on RD, measuring restriction or hindrance perpendicular to the main axis of the bundle, and was therefore summarized as “axon density” component. The second component loaded mostly on MTR, and was thus summarized as “apparent myelin” component.

**Figure 4.**
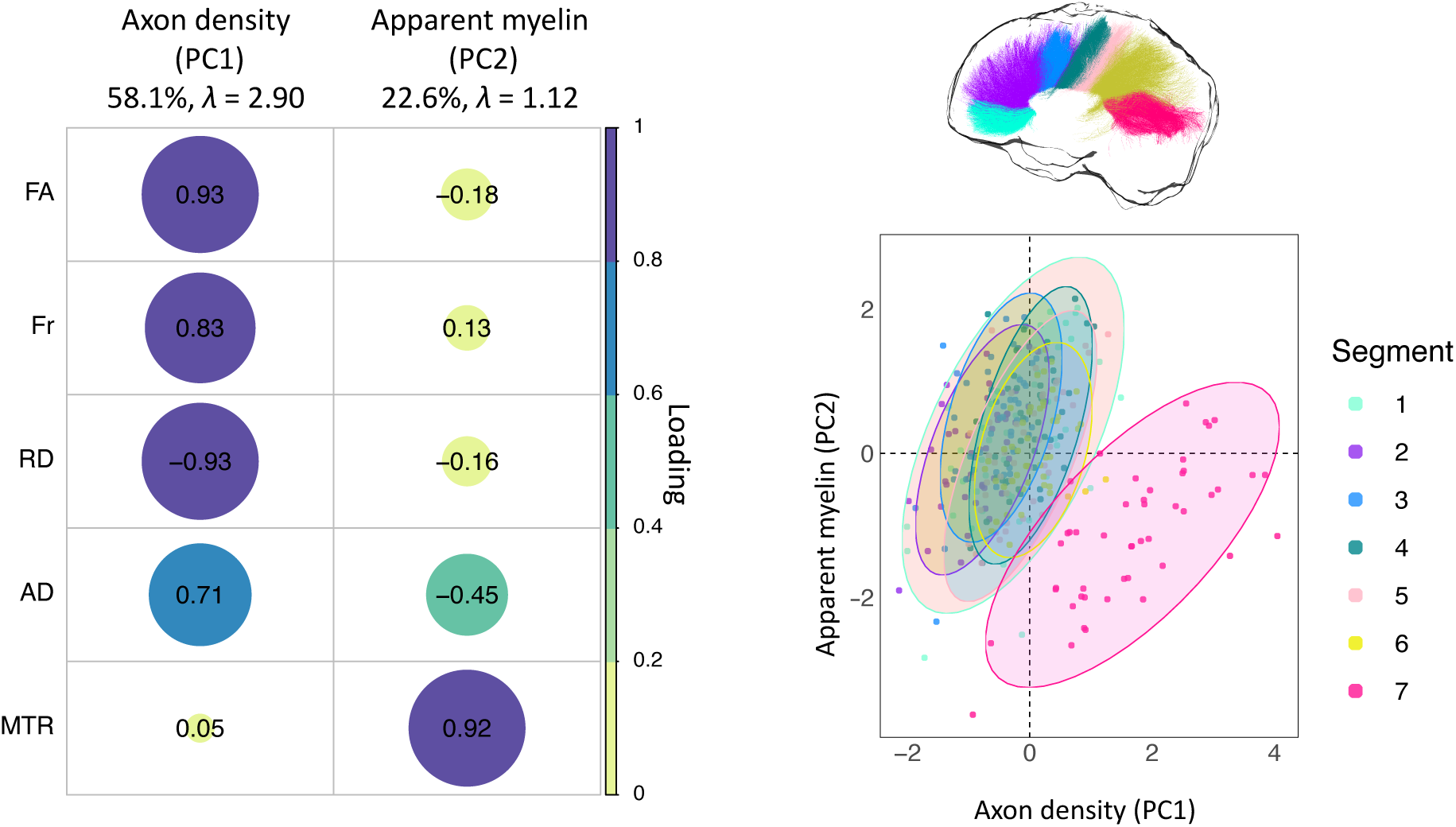
PCA of the microstructure metrics with varimax rotation. Left: Plot summarizing how each variable is accounted for in every principal component. The absolute correlation coefficient is plotted. Color intensity and the size of the circles are proportional to the loading. The final sample size for the PCA was n=25 for the HD group and n=24 for the control group. Right: Segment clustering based on PC1 and PC2. The horizontal axis shows increasing restriction or hindrance perpendicular to the main axis of the bundles. The vertical axis represents an increase in apparent myelin. Each point represents one subject. Concentration ellipsoids cover 95% confidence around the mean. Segment 7 appears to encompass most of the data variability.

### Premanifest patients present alterations in callosal apparent myelin but not axon density

#### Assessment of group differences in axon density

Age was negatively associated with axon density scores (r = -0.301, p < 0.001), and included in the final model assessing the effect of group and segment on axon density scores, with age as covariate.

The effect of group was not significant [F(1, 312) = 1.677, p = 0.196], however a main effect of segment was detected [F(6, 312) = 84.671, p < 0.001] (Fig. 4), together with a main effect of age [F(1, 312) = 34.116, p < 0.001] (Fig. 4). The Group × Segment interaction was not significant [F(6, 312) = 0.531, p = 0.784]. Overall, age was negatively associated with scores on this component; additionally, microstructure in the posterior segments of the CC was associated with higher axon density scores, compared to anterior ones [adjusted means: CC1 = -0.270; CC2 = -0.822; CC3 = -0.546; CC4 = -0.001; CC5 = -0.144; CC6 = 0.083; CC7 = 1.753].

#### Assessment of group differences in the apparent myelin

Age and ICV were correlated with scores on the apparent myelin component (age: r = -0.301, p < 0.001; ICV: r = -0.332, p < 0.001), thus the final model assessed the main effects of group and segment, and age-by-group and a group-by-segment interactions, with age as covariate.

There were no main effects of group [F(1, 312) = 2.353, p = 0.126] or ICV [F(1, 312) = 1.875, p = 0.172]. However, significant main effects of age [F(1,312) = 45.07, p < 0.001] and segment [F(1, 312) = 19.899, p < 0.001] were detected. Overall, scores on this component were lower in segment 7 of the CC and in older participants (Fig. 4).

Crucially, a significant interaction was detected between segment and group [F(6, 312) = 2.238, p = 0.040], indicating that the effect of group was different for different callosal segments. Therefore, slopes of the effect of group on apparent myelin scores for each segment, while controlling for the effect of age, were investigated with a simple moderation analysis using the PROCESS toolbox for SPSS ^78^, to better understand this interaction.

This analysis revealed that patients presented significantly higher apparent myelin compared to controls in segment 1 (p = 0.016), and significantly lower scores in segment 7 (p = 0.0343). Overall, scores on the apparent myelin component for the patient group were higher than controls in the more anterior portions of the CC but lower in the posterior portions (segment1: β = 0.56, t = 2.41, p = 0.016; segment 2: β = 0.25, t = 1.08, p = 0.27; segment 3: β = 0.014, t = 0.06, p = 0.95; segment 4: β = 0.2098, t = 0.90, p = 0.36; segment 5: β = 0.44, t = 1.89, p = 0.058; segment 6: β = -0.028, t = -0.12, p = 0.899; β = -0.5, t = -2.12, p = 0.034) (Fig. 5).

**Figure 5.**
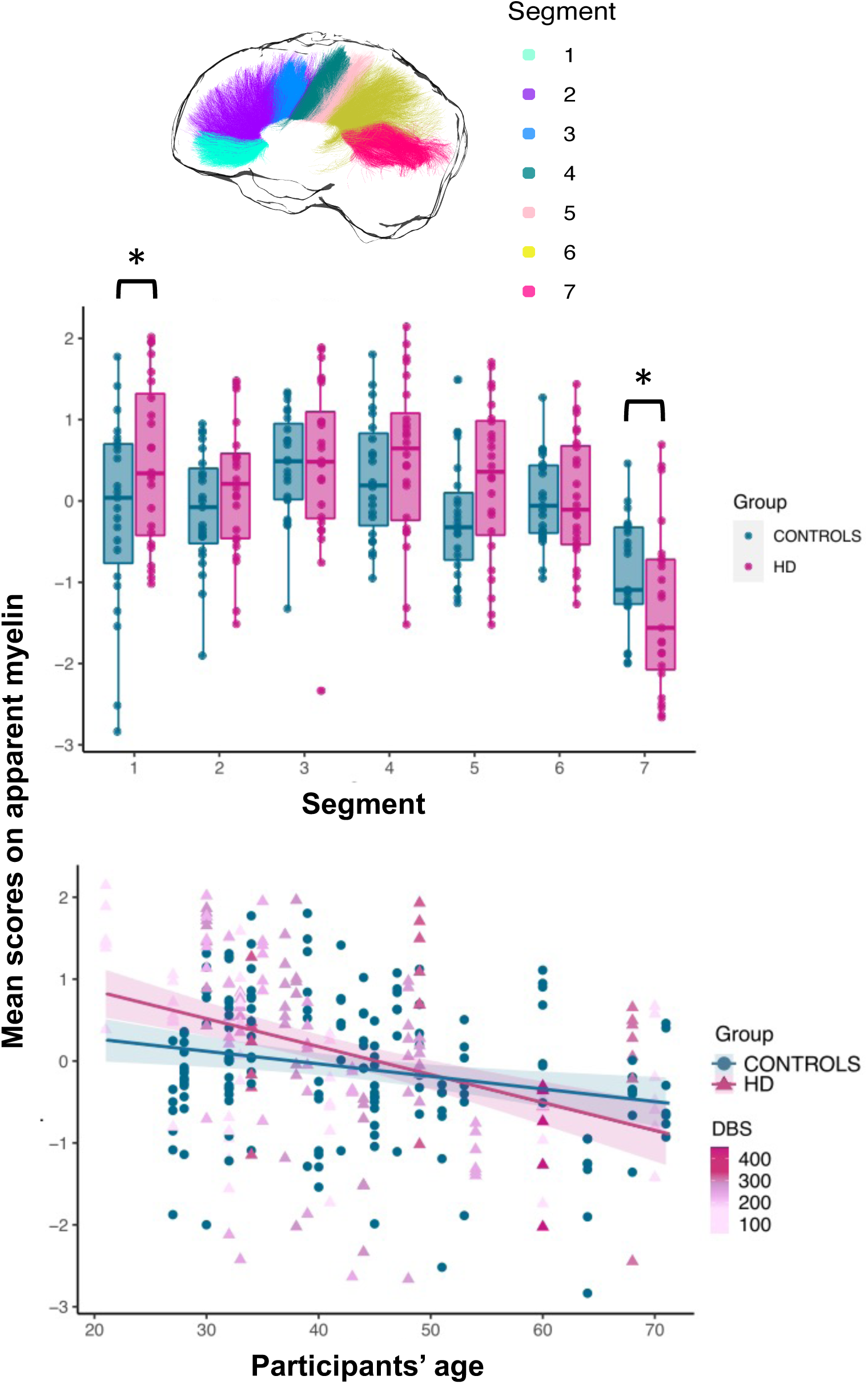
Callosal apparent myelin: patient-control differences across callosal segments (top), and relationship between age and inter-individual variability in apparent myelin (bottom). A group-by-segment interaction effect (p = 0.04) was observed for callosal apparent myelin, indicating that the effect of group was different for different callosal segments. Patients presented significantly higher apparent myelin compared to controls in segment 1 (p = 0.016), and significantly lower in segment 7 (p = 0.034). Overall, scores on the apparent myelin component for the patient group were higher than controls in the more anterior portions of the CC but lower in posterior portions. Additionally, a significant interaction effect between group and age indicated that, while older HD patients presented significantly lower apparent myelin than age-matched controls, the opposite was true for younger HD patients. * p < 0.05, ** p < 0.01, *** p < 0.001, Bonferroni-corrected.

As a post-hoc, exploratory analysis, the impact of partial volume artifacts on apparent myelin differences between patients and controls was assessed. The fractional volume of free water in each voxel was estimated from the diffusion data to produce a free-water signal fraction (FWF) map. The overall mean FWF was then calculated, as described above for the other metrics assessed. Finally, an ANCOVA was run to assess group differences in apparent myelin across the different segments, controlling for FWF. Specifically, the main effects of group and segment and their interaction effect were examined, with age, ICV and FWF as covariates. Age-by-group and group-by-FWF interactions were included in the model because of violation of the homogeneity of regression slopes assumption.

Consistent with the main analysis, a significant main effect of age [F(1, 300) = 56.08, p < 0.001] and segment [F(1, 300) = 22.89, p < 0.001] and a significant interaction effect between segment and group [F(1,300) = 3.2, p = 0.005] were detected. The interaction between group and age [F(1, 300) = 8.736, p = 0.003] was now significant, indicating that while scores on this component are lower than age-matched controls in older patients, the opposite was true for younger patients. Finally, a significant main effect of group [F(1, 300) = 13.042, p < 0.001], and FWF [F(1, 300) = 13.32, p < 0.001], and a significant interaction effect between group and FWF [F(1, 300) = 19.262, p < 0.001], were detected.

#### Apparent callosal myelin is associated with CAG repeat length but not with cognitive performance or disease burden

Spearman correlation coefficients and associated p-values for the correlations of apparent callosal myelin with composite cognitive scores, CAG repeat length and DBS are reported in Table 4. Trends for positive associations were detected between composite cognitive scores and apparent myelin in all segments, except for segment 7. However, these associations were no longer significant after multiple comparison correction. Apparent myelin was positively correlated with CAG repeat length in segment 1 (r = 0.641, p = 0.002), segment 2 (r = 0.717, p = 0.001), segment 3 (r = 0.549, p = 0.012), segment 4 (r = 0.549, p = 0.012), segment 5 (r = 0.525, p = 0.018), and segment 6 (r = 0.513, p = 0.021). After Bonferroni correction the relationship remained significant in segments 1 (p = 0.014), 2 (p = 0.007) and 4 (p = 0.007) (Fig. 6). Partial correlations were carried out to explore the relationships between apparent myelin and CAG repeat length independently of ICV and disease burden. Even stronger positive associations were now detected; interestingly, the association was now significant also in segment 7, before correction (segment 1: r = 0.763, p = 0.001, corrected p = 0.007; segment 2: r = 0.879, p < 0.001, corrected p < 0.001; segment 3: r = 0.841, p < 0.001, corrected p < 0.001; segment 4: r = 0.83, p < 0.001, corrected p < 0.001; segment 5: r = 0.745, p = 0.001, corrected p = 0.007; segment 6: r = 0.864, p < 0.001, corrected p < 0.001; segment 7: r = 0.5, p = 0.048, corrected p = 0.336) (Fig. 5). No significant associations were detected between apparent myelin scores in each of the 7 callosal segments and DBS.

**Figure 6.**
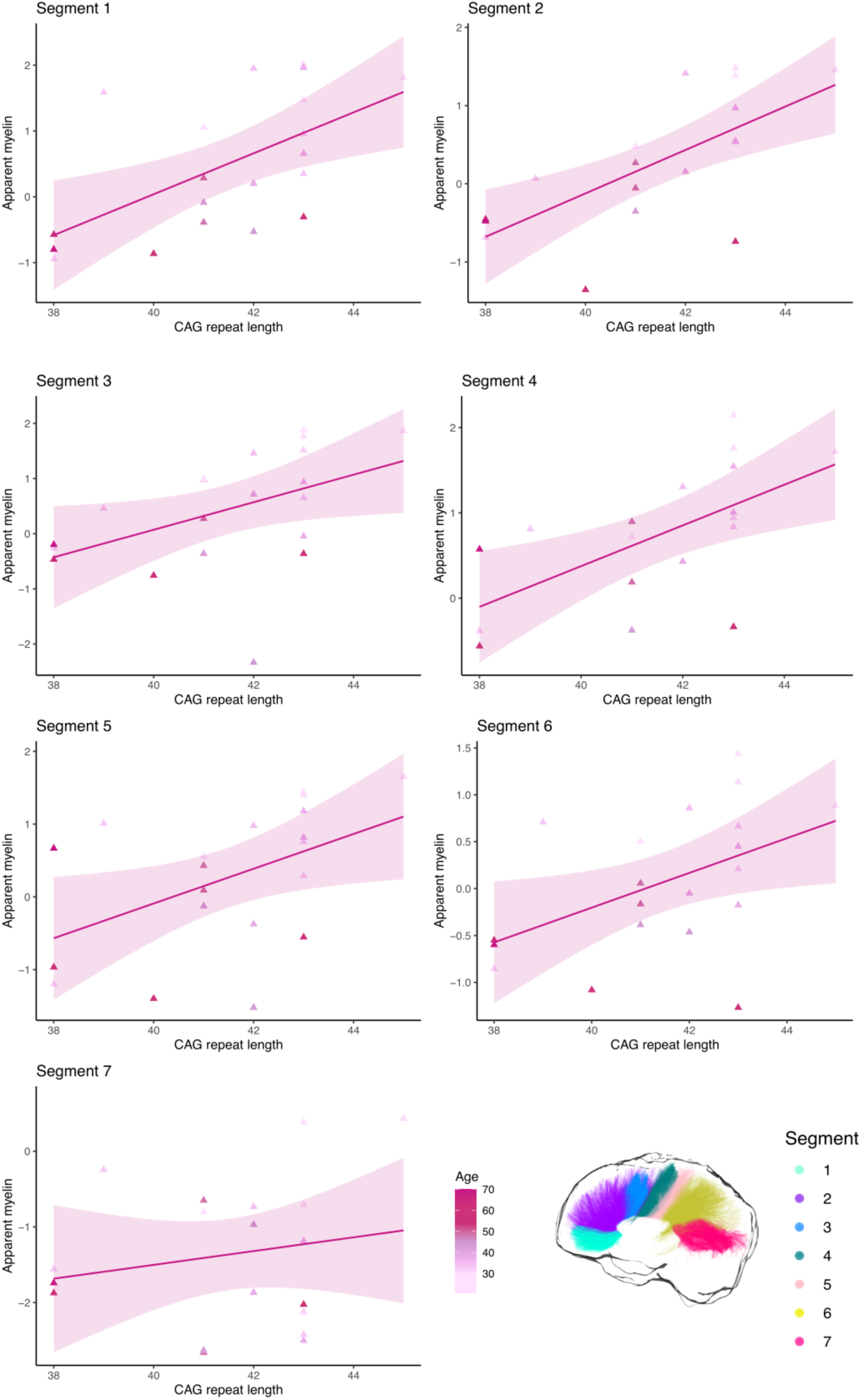
Relationship between apparent myelin in each callosal segment and CAG repeat length in patients.

**Table 4.** Correlations of apparent myelin scores with cognitive component scores, CAG repeat-length and DBS.

| **Apparent myelin** | **Composite cognitive scores** |
| --- | --- |
| **Segment 1** | $r = 0.527$ ($p = 0.032$, corrected $p = 0.211$) |
| **Segment 2** | $r = 0.559$ ($p = 0.023$, corrected $p = 0.141$) |
| **Segment 3** | $r = 0.491$ ($p = 0.042$, corrected $p = 0.282$) |
| **Segment 4** | $r = 0.494$ ($p = 0.054$, corrected $p = 0.351$) |
| **Segment 5** | $r = 0.451$ ($p = 0.073$, **corrected $p = 0.049$**) |
| **Segment 6** | $r = 0.323$ ($p = 0.03$, corrected $p = 0.213$) |
| **Segment 7** | $r = -0.098$ ($p = 0.71$, corrected $p = 1$) |
|  | **CAG repeat length** |
| **Segment 1** | $r = 0.641$ ($p = 0.002$, **corrected $p = 0.014$**), partial correlation: $r = 0.763$ ($p = 0.001$, **corrected $p = 0.007$**) |
| **Segment 2** | $r = 0.717$ ($p = 0.001$, **corrected $p = 0.007$**), partial correlation: $r = 0.879$ ($p < 0.001$, **corrected $p < 0.001$**) |
| **Segment 3** | $r = 0.549$ ($p = 0.012$, corrected $p = 0.084$), partial correlation: $r = 0.841$ ($p < 0.001$, **corrected $p < 0.001$**) |
| **Segment 4** | $r = 0.71$ ($p = 0.001$, **corrected $p = 0.007$**), partial correlation: $r = 0.831$ ($p < 0.001$, **corrected $p < 0.001$**) |
| **Segment 5** | $r = 0.525$ ($p = 0.018$, corrected $p = 0.126$), partial correlation: $r = 0.745$ ($p = 0.001$, **corrected $p = 0.007$**) |
| **Segment 6** | $r = 0.513$ ($p = 0.021$, corrected $p = 0.147$), partial correlation: $r = 0.864$ ($p < 0.001$, **corrected $p < 0.001$**) |
| **Segment 7** | $r = 0.107$ ($p = 0.663$, corrected $p = 1$), partial correlation: $r = 0.5$ ($p = 0.048$, corrected $p = 0.336$) |
|  | **DBS** |
| **Segment 1** | $r = -0.04$ ($p = 0.853$, corrected $p = 1$) |
| **Segment 2** | $r = 0.08$ ($p = 0.697$, corrected $p = 1$) |
| **Segment 3** | $r = 0.003$ ($p = 0.986$, corrected $p = 1$) |
| **Segment 4** | $r = 0.071$ ($p = 0.739$, corrected $p = 1$) |
| **Segment 5** | $r = 0.048$ ($p = 0.824$, corrected $p = 1$) |
| **Segment 6** | $r = -0.12$ ($p = 0.642$, corrected $p = 1$) |
| **Segment 7** | $r = -0.09$ ($p = 0.662$, corrected $p = 1$) |

*Correlation coefficients that were significant after Bonferroni correction are highlighted in bold. Trends, defined as correlations significant at the uncorrected level, are highlighted in italics.*

#### Whole-brain analysis with TBCA reveals WM microstructure alterations in the posterior CC, the left CST and the right fronto-striatal projections

Fig. 7A shows the TBCA results. Consistent with the PCA results, a significant reduction in MTR in the patient group was detected, compared to controls, in the posterior portion of the CC [cluster mass (∑ t-score) = 1530, p < 0.001 (uncorrected), p = 0.030 (FWE-corrected)]. Furthermore, a significant increase in FR along most of the left CST was found in patients [cluster mass (∑ t-score) = 1004, p < 0.001 (uncorrected), p = 0.030 (FWE-corrected)]. Finally, right-lateralized clusters of significantly higher FA in the patient group were identified in the fronto-striatal projections [cluster mass (∑ t-score) = 956, p < 0.001 (uncorrected), p = 0.03 (FWE-corrected)].

**Figure 7.**
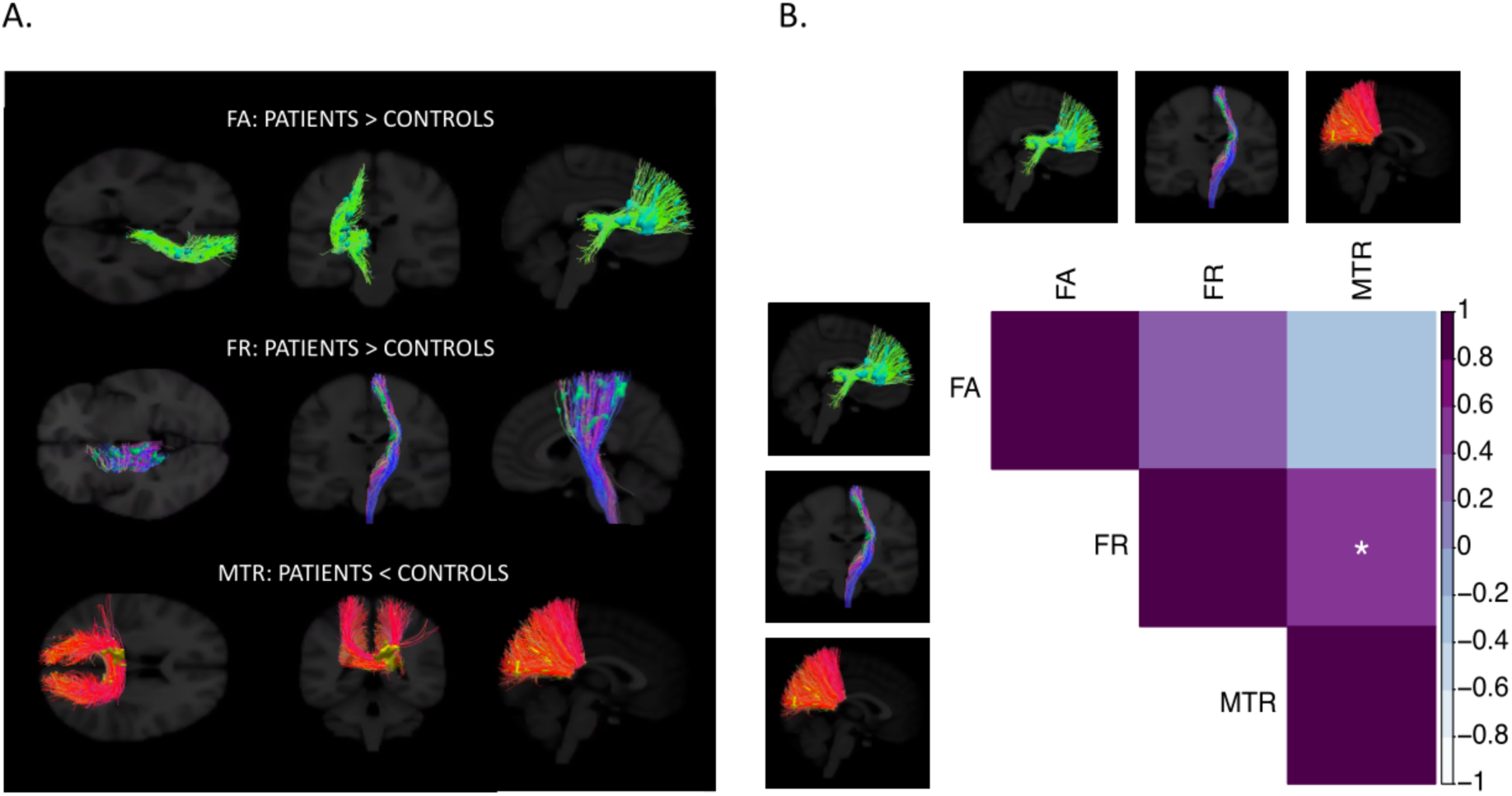
Results of the cluster-analysis obtained with TBCA between patients and controls (A) and Spearman correlations between significant TBCA clusters in patients (B) * p < 0.05, ** p < 0.01, *** p < 0.001, Bonferroni-corrected.

Fig. 7B plots the relationship between significant microstructure clusters as detected with TBCA for patients. FR in the CST was significantly associated with MTR in the posterior CC (r = 0.498, p = 0.011, corrected p = 0.033), but not with FA in the right fronto-striatal projections (r = 0.328, p = 0.110, corrected p = 0.327). Additionally, MTR was not associated with FA (r = -0.218, p = 0.294, corrected p = 0.882).

## Discussion

We carried out a comprehensive tractometry analysis ^16–18^ of regional changes across the CC in premanifest HD. By exploiting the ultra-strong magnetic field gradients of the Connectom scanner ^8,9^, it was possible to better tease apart alterations in myelin content from alterations in axon microstructure ^10^.

Lower apparent myelin, but not axon density, was detected in the callosal isthmus in patients with both analytical approaches. Previous DTI studies have reported microstructure changes in this callosal region in premanifest patients ^82,90^. Here, the combination of standard DTI metrics with MTR and FR afforded a more biologically meaningful interpretation of microstructural changes. These results replicate evidence from a previous study carried out by our group at 7 Tesla ^7^ and are in accord with the Demyelination Hypothesis, which argues that early myelinated fibres are more susceptible to myelin disorder in the disease ^21^.

Interestingly, patients presented significantly higher apparent myelin than controls in the callosal rostrum and, overall, apparent myelin was higher in patients than controls in the more anterior portions of the CC. Additionally, a positive association was detected between apparent myelin and CAG size in patients, indicating that myelin alterations are directly linked to the disease mutation. Finally, a significant interaction effect was detected between group and age on apparent myelin, suggesting that while myelin content in this tract is higher in younger patients, the opposite is true for older patients, which likely present increased disease burden.

Based on these findings, it is possible that, at least early on in disease progression, the HD mutation is associated with excessive, rather than reduced, myelin production. This might be caused by a pathological increase in myelin-producing, iron-rich oligodendrocytes. In accord with this proposal, previous evidence has suggested that HD gene expression may influence brain cell densities from early in the life of gene carriers ^22^. Additionally, this explanation agrees with findings from neuropathology showing increased density of oligodendrocytes in the brain of premanifest patients ^23^; furthermore, *mHTT* directly alters the proliferation property of cultured oligodendrocyte precursor cells (OPCs), with the degree of cell proliferation of OPCs increasing with pathological severity and increasing CAG repeat length ^25^. Finally, recent evidence from the cross-sectional HD Young Adult Study demonstrated increased R1 and R2* values, suggestive of either increased iron or increased myelin, in the putamen, globus pallidum and external capsule of HD patients more than 20 years away from clinical onset ^91^.

While earlier in disease progression the disease mutation may be associated with increased myelin, such increases in myelin content may lead to detrimental effects later in the disease due to oxidative stress ^21,92,93^. Critically, lower apparent myelin in the most posterior areas, through which fibres from the visual system transverse, suggests that these regions may be the first to be affected, in agreement with previous evidence ^21,33,94^. The visual system is functionally critical early in life, with myelination occurring early and progressing rapidly ^95^. Additionally, this system is highly dynamic and is associated with big energetic demands. As metabolic dysfunction and alterations in energetics play important mechanistic roles in HD ^96,97^, these changes may contribute to early myelin impairment in this callosal portion.

Overall, we demonstrate measurable and significant differences in callosal apparent myelin before changes in proxy metrics of axon density can be detected. These changes may reflect early neuronal dysfunction ^98^ or a CAG-driven neurodevelopmental component to the pathogenesis of HD, as a precursor to the more global neurodegeneration process ^25,99–101^. Accordingly, there is increasing evidence that neurodevelopment is affected in HD ^99,102^ and that such developmental elements of HD are independent of ongoing neurodegeneration ^91^. Nevertheless, as the present study was not designed to detect HD-associated developmental changes, future studies following young premanifest subjects longitudinally should address the possibility of toxic myelin levels because of pathological CAG repeats size.

With TBCA, clusters of significantly higher FA were detected in the patient group in the right fronto-striatal projections. Though neurodegenerative disorders have normally been associated with lower FA in major WM pathways, attributed to WM degeneration, demyelination, reduced gliosis or axonal damage as a result of GM loss ^103,104^, it is possible that selective degeneration of specific WM tracts resulted here in higher anisotropy values and a paradoxical increase in microstructural organization^105^. This suggests that WM degeneration in this area is already present at the premanifest stage of the disease.

Importantly, significantly higher FR along most of the left CST was also detected with TBCA. This tract is composed of descending WM fibres, with half of them arising from the primary motor cortex, and is anatomically linked to the basal ganglia ^106,107^. From a functional point of view, the CST conducts motor impulses from the brain to the spinal cord, and plays an essential role in voluntary movement ^106,107^. Though the hallmark symptom of HD concerns involuntary choreic movements ^108^, alterations in voluntary movement are also present in premanifest patients ^109^, thus suggesting that alterations in this tract may play an important role in the disease. Crucially, this is the first time that alterations in this measure have been detected in premanifest patients, pointing to the potential of FR as *in-vivo* MRI marker of premanifest neural changes.

Previous studies have demonstrated lower WM volume in the internal capsule of manifest patients ^110,111^. Accordingly, the elevated FR detected in this study might reflect the loss of non-neuronal cells, in turn leading to axons being pushed together ^112^. Alternatively, such a result might reflect axonal swelling ^113^. Consistent with this suggestion, previous evidence demonstrated higher iron levels in the left CST of premanifest patients ^91,114^, interpreted as indicating an homeostatic increase in oligodendrocytes to repair myelin damage. In turn, myelin damage leads to axon swelling ^115^. It might also be that fibre bundles develop differently because of the genetic mutation, and this is consistent with evidence of morphological alterations in the neurons of HD mice, which present smaller diameter dendritic shafts, smaller somatic cross-sectional areas, and decreased diameter of the dendritic fields ^116^. Finally, higher FR might reflect the presence of a process of reorganization and compensatory pruning of axons in WM, such as pathologically-driven reduced collateral branching or morphological alterations of individual axons. Consistent with this suggestion, evidence has shown increased coherence of axonal organization in premanifest patients, as suggested by a smaller orientation dispersion index (OD), in tracts surrounding the basal ganglia and in the internal and external capsule ^117^.

The finding of higher FR in the left CST is consistent with the leftward-biased GM loss demonstrated in the striatum of patients ^118^ and with the leftward asymmetry of brain iron in aging and motor disorders ^119,120^. Nevertheless, future studies are needed to determine whether this is an important finding to understand disease pathology. For example, future studies could investigate the longitudinal evolution of changes in FR in patients.

To date, only one other study has used extensive microstructural measures in premanifest HD^91^. Nevertheless, such measurements are essential for understanding the trajectory of myelin alterations across the disease course, which is expected to vary as disease processes change ^91,121^. Notably, though much of our understanding of HD pathology will increasingly rely on advanced neuroimaging techniques, it is important to remember and address the shortcomings of these approaches. For example, MTR is influenced by a complex combination of biological factors (including T_1_), making it difficult to separate the effects of reduced macromolecular density because of demyelination and/or axonal loss, or increased water because of oedema and/or inflammation ^122–125^. Future investigations may benefit from utilising quantitative magnetization transfer imaging techniques ^126^ to assess myelin alterations in the premanifest disease stage. Additionally, because of the way FR is computed, a change in T_2_ relaxation (for example because of altered tissue water or myelin content) may be erroneously interpreted as a difference in FR. Thus, future studies are needed to clarify the neurobiological underpinning of our finding, for example by investigating disease-associated changes in volume and axon diameter distribution in the CST.

Notwithstanding the above limitations, findings from this work highlight the fundamental importance of gaining an enhanced understanding of the mechanisms underlying WM abnormalities in HD. Crucially, our results suggest that myelin alterations in the disease may reflect CAG-driven neurodevelopmental, rather than neurodegenerative, changes and that expanding intervention strategies to include oligodendroglial targets ^28^ directly targeting WM pathology may be beneficial for HD.

## Availability of data and materials

The data analysed during the current study and the respective analysis scripts are available from the corresponding author on reasonable request.

## Funding and acknowledgements

The present research was funded by a Wellcome Trust PhD studentship to CC (ref: 204005/Z/16/Z); DKJ was supported by a New Investigator Award (to DKJ) from the Wellcome Trust (ref: 096646/Z/11/Z) and a Strategic Award from the Wellcome Trust (ref: 104943/Z/14/Z). We thank Dr Slawomir Kusmia and Dr Elena Kleban for their support with the project.

## Competing interests

The authors report no competing interests.

## Abbreviations

AD: axial diffusivity
ANCOVA: analysis of covariance
CAG: cytosine, adenine and guanine
CC: corpus callosum
CHARMED: Composite Hindered and Restricted Model of Diffusion
CST: cortico-spinal tract
DBS: disease burden score
DCL: diagnostic confidence level
DT MRI: diffusion tensor MRI
EV: explanation variable
FA: fractional anisotropy
FAS: functional assessment score
FDR: false discovery rate
fODF: fiber orientation density function
FoV: field of view
FR: restricted fraction
FEW: family wise error
FWF: free-water fraction
HTT: huntingtin
ICV: intracranial volume
KMO: Kaiser=Meyer-Olkin
MD: mean diffusivity
*mHTT*: mutant huntingtin
MoCA: Montreal Cognitive Assessment
MPRAGE: magnetization prepared – rapid gradient echo
MT: magnetization transfer
MT-w: MT-weighted
MTI: magnetization transfer imaging
MTR: magnetization transfer ratio
OPC: oligodendrocyte precursor cells
PCA: principal component analysis
PEBL: Psychology Experiment Building Language
RD: radial diffusivity
ROI: region of interest
SD: standard deviation
TBCA: tract-based cluster analysis
TFC: total functional score
TMS: total motor score
TOPF-UK: Test of Premorbid Functioning – UK Version
TOPF-UK FSIQ: TOPF-UK full scale IQ
UHDRS: Unified Huntington’s Disease Rating Scale
WAIS-R: Wechsler Adult Intelligence Scale-Revised
WM: white matter

## Notes

### Competing Interest Statement

The authors have declared no competing interest.

